# Characterizing interactions of ER resident proteins in situ through the YST-PPI method

**DOI:** 10.1101/2024.05.21.594841

**Authors:** Xian Fan, Huahua He, Ting Wang, Pan Xu, Faying Zhang, Shantong Hu, Yueli Yun, Meng Mei, Guimin Zhang, Li Yi

## Abstract

The mutual interactions of ER resident proteins in the ER maintain its functions, prompting the protein folding, modification, and transportation. Here, a new method, named YST-PPI (YESS-based Split fast TEV protease System for Protein-protein Interaction) was developed, targeting the characterization of protein interactions in ER. YST-PPI method integrated the YESS system, split-TEV technology and endoplasmic reticulum retention signal peptide (ERS) to provide an effective strategy for studying ER in situ PPIs in a fast and quantitative manner. The interactions among 15 ER resident proteins of *S. cerevisiae* were explored using the YST-PPI system, and their interaction network map was constructed, in which more than 74 interacting resident protein pairs were identified. Our studies also showed that Lhs1p plays a critical role in regulating the interactions of most of the ER resident proteins, expect the Sil1p, indicating its potential role in controlling the ER molecular chaperones. Moreover, the mutual interaction revealed by our studies further confirmed that the ER resident proteins perform their functions in a synergetic way and multimer complex might be formed during the process.

## 1 INTRODUCTION

The correct protein folding is very critical to human cells, whose dysfunction will cause serious diseases. In eukaryotic cells, the ER and Golgi apparatus are major organelles for protein folding, post-translational modification and transportation ^[1]^. Generally, these process happened in ER are under the assistance by molecular chaperones and folding enzymes ^[2]^, among which a group of proteins, called ER resident proteins, play important roles in maintaining ER homeostasis, such as Bip and Sil1p in human cells, and Lhs1p and Kar2p in yeast cells, etc ^[3]^.

To maintain their residence in ER and Golgi, the ER resident proteins share a similar xxH/KDEL sequence at C-terminal, named ER retention sequence (ERS) ^[4]^, to mediate their interaction with ERS receptors ^[5]^. The main ERS in human cells is KDEL, which can bind to KDELR1, KDEL2 and KDELR3 receptors to achieve retention effect ^[6]^. Comparably, the main ERS in *S. cerevisiae* is HDEL, which binds to the Erd1p or Erd2p receptors ^[5]^. It is worthy to pointing out that ERS is not the only mechanism to retain ER resident proteins. For example, Ero1p, an important protein together with Pdi1p involved in protein disulfide bond formation in ER, does not have C-terminal HDEL ^[7]^.

All ER resident proteins exist in ER to carry out different but collaborative chaperone functions to prompt maturation of newly synthesized protein ^[8]^. Their dysfunction or absence will cause the accumulation of misfolded proteins in ER ^[9]^, which might disrupt ER homeostasis, leading to apoptosis and even some related prevalent diseases, such as diabetes, neurodegeneration and liver and heart diseases ^[10]^. Because of the co-existence and collaborative functions of ER resident proteins, there raises an interesting question that if mutual interactions exist to maintain the correct ER functions. Unfortunately, very limited studies of PPIs among ER resident proteins have been explored due to the lack of ER in situ methods. The general protein interaction databases BioGRID (4.4) ^[11]^ and STRING ^[12]^ provide some ER PPIs information, while most of the data were extracted from in *vitro* assays instead of under physiological conditions ^[13]^.

Studying PPIs in human ER is still very challenging due to the complexity of human cells. At the same time, *S. cerevisiae*, as a eukaryotic model organism, can also be genetically manipulated with well-studied biological background ^[14]^. It has recently been reported that 47% of *S. cerevisiae* genes can be replaced by human genes ^[15]^, further indicating that it can provide a theoretical basis for human life science research. Therefore, studying the interactions of ER resident proteins in *S. cerevisiae* can provide insights of the cooperative modes in their functional processes, thus expanding the industrial and pharmaceutical applications of yeast cells and guiding further understanding of protein folding and function in human cells.

Here, we developed the YST-PPI system, which integrates the YESS system ^[16]^, split-TEV method ^[17]^ and ERS ^[18]^, for fast and quantitative characterization of PPIs in yeast ER (**Figure 1**). Using the YST-PPI system, the interacting network of 15 ER resident proteins in *S. cerevisiae*, which contain the conserved DEL sequence at C-terminus, was analyzed. We identified more than 74 interacting pairs of ER-resident proteins with a normalized interacting score higher than 10%. In addition, the potential networking regulatory roles of Lhs1p in protein folding was also explored, which laid the foundation for in-depth analysis of the synergistic interaction of ER resident proteins.

**FIGURE 1.**
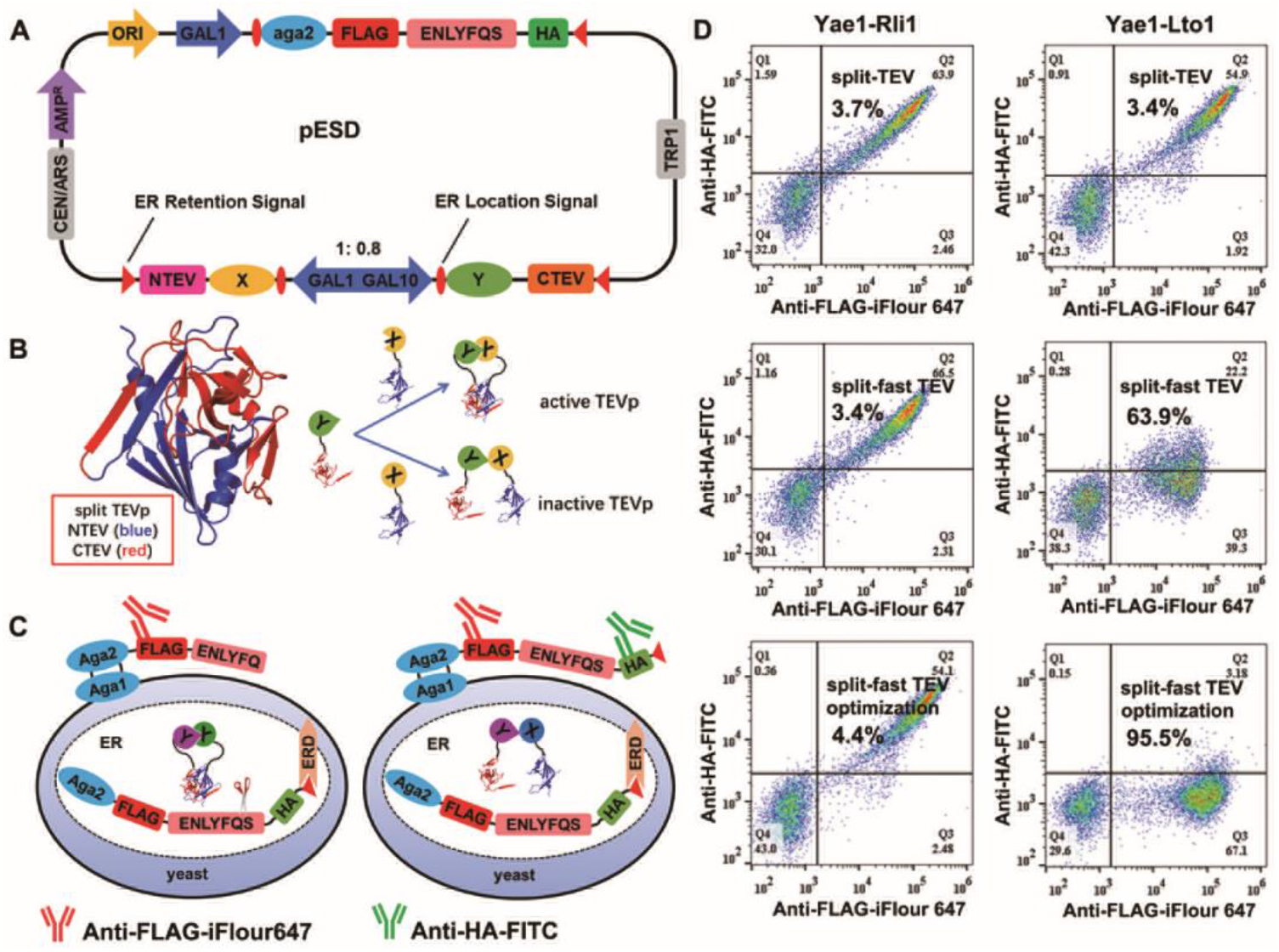
The principle of YST-PPI system. A, the schematic diagram of YST-PPI plasmid with three expression frames, in which the GAL1 promoter is fused with aga2-FLAG-ENLYFQS-HA-ERS substrate complex, and the GAL1-GAL10 bidirectional promoter guides X-NTEV-ERS and Y-CTEV-ERS target protein complex. B, the schematic diagram of split-TEV interaction with target protein (PDB: 1LVB ^[22]^). C, the schematic diagram of the substrate complex on the surface of S. cerevisiae cells. When the prey and bait proteins interact with each other, the substrate complex is cleaved, followed by display on yeast cell surface, in which the FLAG epitope tag is labeled by fluorescent antibody. When the prey and bait proteins do not interact with each other, the substrate complex is not cleaved, followed by display of intact substrate on yeast cell surface, in which both FLAG and HA epitope tags are labeled by different fluorescent antibodies. ERD, ERS receptor Erd1p or Erd2p. D, FACS results of positive control (Yae1p-Lto1p) and negative control (Yae1p-Rli1p) under the condition of the stronger ERS (FEHDEL) at C-terminal of both prey and bait complex, using 41AA as Linker1 and 2G4S as Linker2, in which the split-TEV and split-fast TEV are used for the assay, respectively. The “optimization” shows the optimal interaction of positive and negative controls after optimizing ERS and Linker based on split-fast TEV.

## 2 MATERIALS AND METHODS

### 2.1 Vector construction

In this manuscript, the pESD plasmid containing three expression frames was modified to detect interaction of ER resident proteins ^[16]^. The bidirectional promoter GAL10/GAL1 has similar activity and controls the expression of the two target protein complexes (prey protein X-NTEV-ERS and bait protein Y-CTEV-ERS) (**Figure 1A**), respectively. A single GAL1 promoter regulates the expression of a substrate complex (Aga2-FLAG-ENLYFQ↓G-HA-ERS) (**Figure 1A**), and Aga2 can bind to Aga1 on the surface of *S. cerevisiae*, thereby displaying the substrate complex on the cell surface.

### 2.2 Yeast strain modification

*S. cerevisiae* EBY100 (*URA* ^+^, *leu* ^-^, *trp* ^-^) is used for yeast surface display. The upstream and downstream *lhs1* gene fragments of about 700bp in the genome of EBY100 were amplified, and then connected to both ends of the *G418* resistance gene for recombination into the genome of EBY100 strain. Finally, *lhs1* gene deletion strain EBY100(Δ*lhs1*) was identified by G418 antibiotics resistance screening and PCR identification (**Figure S1**).

For the EBY100(+*lhs1*) strain generation, the expression frames of *lhs1* and *Cre* ^[19]^ were inserted into pGAPZB plasmid, respectively. After linearization, the gene fragment -*lhs1* up*-* G*AP-lhs1-CYC1-Loxp71-GAL1-Cre-CYC1-Zeocin-Loxp66-lhs1* down-was obtained, in which two homologous arms (*lhs1* up and *lhs1* down) were used for recombination into the genome of EBY100 (Δ*lhs1*) strain. The transformants were screened by using plates containing *Zeocin* and *G418* antibiotics. The strains that grew on the *Zeocin* resistant plate but not on the *G418* resistant plate were selected as successfully integrated strains. Subsequently, the Cre expression is induced to knock out the gene fragment (-GAL1-Cre-CYC1-*Zeocin* -). The galactose induced GAL1 promoter to activate Cre enzyme expression, which can be used to identify specific gene sequences Loxp71 and Loxp66 to knock out the gene fragments between ^[20, 21]^. The final integrated EBY100(+*lhs1*) strain was verified by antibiotics resistance screening and PCR identification (**Figure S2**). The transcription level of the *lhs1* gene in the EBY100, EBY100(Δ*lhs1*), and EBY100(+*lhs1*) strains were characterized and *lhs1* transcription level is about 10 times higher than that of the EBY100 strain (**Figure S2C)**.

### 2.3 FACS analysis

The above constructed pESD plasmids were transformed into EBY100, EBY100(Δ*lhs1*), and EBY100(+*lhs1*) strains, followed by growing on SD plate, respectively. When the single colony grew in SD-Glu (glucose, CAA, YNB) to OD600 of 4-6, some cells were transferred to SD-Gal (galactose, CAA, YNB) medium with an initial OD600 of 0.8. The expression levels of the target protein and substrate complexes reached the highest after 12-16h induction. Subsequently, yeast cells displayed with the substrate complex were labeled with iFluor 647 conjugated anti-FLAG and FITC conjugated anti-HA antibodies (GenScript, Nanjing, China). The fluorescence signals of labeled cells were quantitated with Beckman Coulter CytoFLEX Flow Cytometer (Beckman Coulter, Brea, CA, USA) using previously published protocols. The iFluor 647 and FITC fluorescent intensity were detected with the APC channel (660/20 nm BP) and FITC channel (525/40 nm BP), respectively. The normalized interacting score was calculated as:

[Relative strength of protein interaction] = ([Cells exhibiting only iFluor 647 fluorescence signals]/ ([Cells exhibiting only iFluor 647 fluorescence signals] + [Cells exhibiting both iFluor 647 and FITC fluorescence signals]) x 100%

### 2.4 Protein Interaction comparison

The PPIs identified by YESS system were reconfirmed by BioGRID (Biological General Repository for Interaction Datasets) ^[11]^ and STRING (search tool for recurring instances of neighboring genes) ^[12]^ protein interaction databases.

## RESULTS

### 3.1 Development of YST-PPI system

In original YESS system, TEV protease (TEVp) could cleave its substrate (ENLYFQ↓S) in the yeast ER, followed by transporting the cleaved and uncleaved substrates to the yeast surface for quantification ^[16]^. Here, we integrated the split-TEV technology ^[17]^ into the YESS system, developing the YST-PPI system to characterize PPIs in yeast ER. In YST-PPI, the two target proteins, X (prey) and Y (bait), were fused to N-TEV (1-118) and C-TEV (119-242), respectively (**Figure 1A**). When target proteins X and Y interacts, the active TEVp (PDB: 1LVB ^[22]^) can be reassembled by N-TEV and C-TEV to cleave its substrates (**Figure 1B**). Then the cleaved and uncleaved substrates were transported to yeast cell surface, followed by fluorophore conjugated antibody labeling and flow cytometry quantification (**Figure 1C**).

First, we used three well studied yeast proteins, Yae1p, Rli1p, and Lto1p, to validate the YST-PPI system, in which Yae1p has been proven to only interact with Lto1p ^[23]^. Nevertheless, in our initial setting when wt split-TEV was used, no interaction between Yae1p and Lto1p was detected (**Figure 1D**). We speculated that this might be due to the low activity of resembled wt TEVp, as it was reported that only 30%-40% activity can be restored after the reassembly ^[17]^. To improve the sensitivity of the YST-PPI system, we replace the wt TEVp with an engineered TEVp variants, fast TEVp (G79E, T173A, S219V), which showed a 4-time higher proteolytic activity than that of wt TEVp ^[16]^. We kept the same split sites in fast TEVp as its simulated structure is highly similar with that of wt TEVp (**Figure S3**). As expected, the interaction between Yae1p and Lto1p was better detected using split-fast TEV, presented an enhanced interacting score of 63.9%. Under the same condition, the interacting score of the negative control of Yae1p-Rli1p protein pair was still low (3.4%) (**Figure 1D**).

### 3.2 Optimization of YST-PPI system

To further increase the sensitivity of the YST-PPI system, various ERS was fused to the C-terminal ends of target proteins (**Figure S4A**). In our previous studies, we showed that different C-terminal ERS presented different capability to retain the target proteins in yeast ER, with ER retention strength in the order of WEHDEL > FEHDEL > HDEL > None ^[18]^. In principle, stronger ERS could retain the prey and bait proteins in yeast ER longer to enhance their interacting possibility, thus increasing the sensitivity of the YST-PPI system. Based on our evaluation, the interacting score of Yae1p-Lto1p is only 3% when no ERS was used (as shown in NONE in **Figure 4A**). In comparison, when WEHDEL ^[18]^, the strongest ERS, was fused to the C-terminus of both prey and bait protein complexes and the substrate complex, the interacting score of Yae1p-Lto1p was approximately 90% (as shown in WEHDEL in **Figure 2A** ). FEHDEL and HDEL, which are the moderate ERS, led to a 63% and 20% interacting score of Yae1p-Lto1p, respectively. These results are consistent to the retention capabilities of the different ERS.

**FIGURE 2.**
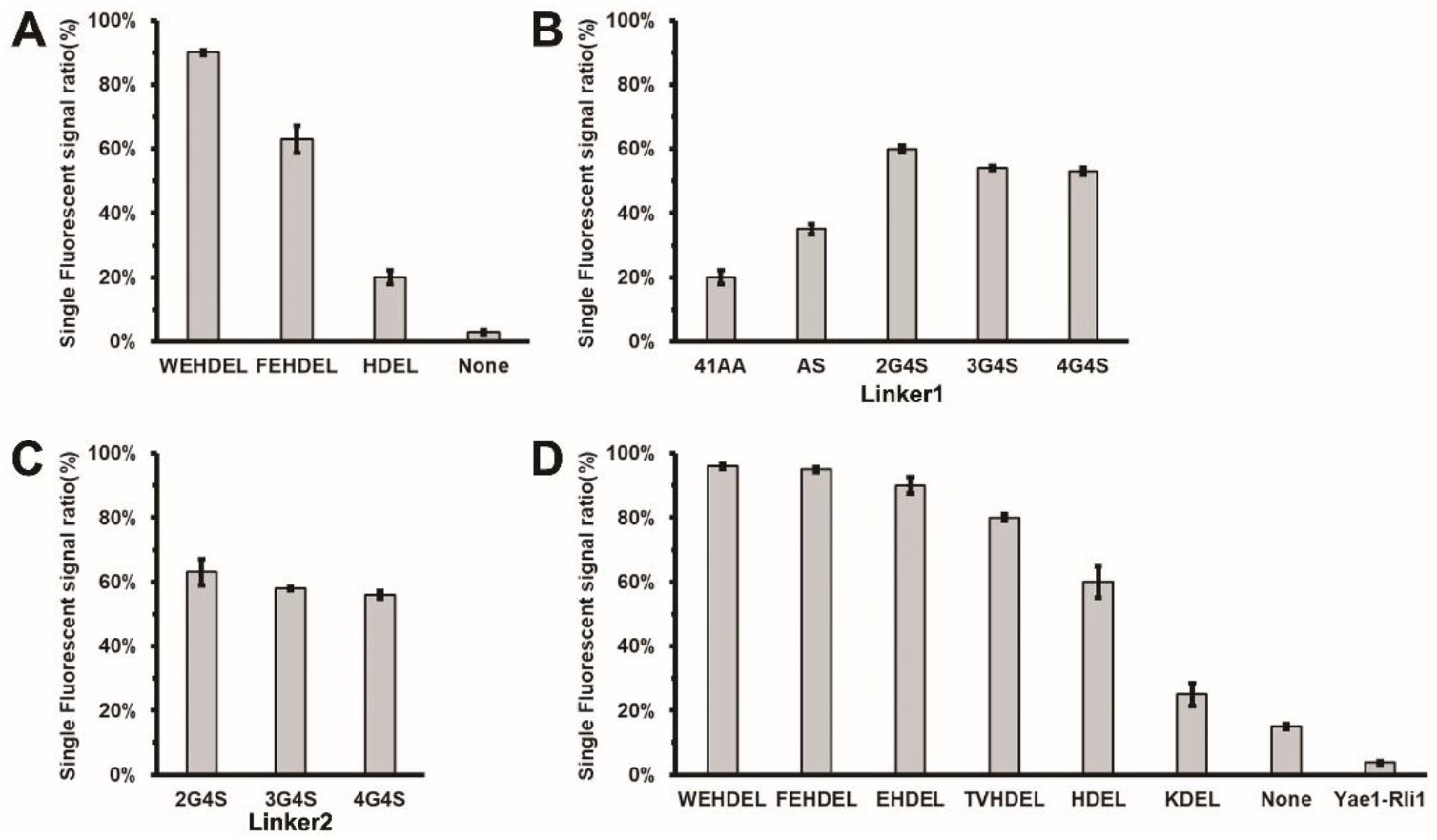
The optimization of YST-PPI system. The Yae1p and Lto1p was fused with N-fast TEV and C-fast TEV, respectively. The Yae1p-Lto1p interacting score was calculated to evaluate the sensitivity of YST-PPI system. A, the effect of different ERS under the condition of using 41AA as Linker1 and 2G4S as Linker2. B, the effect of different Linker1 under the condition of the weak ERS (HDEL) at C-terminal of both prey and bait complex, and using 2G4S as Linker2. C, the effect of different Linker2 under the condition of the ERS (FEHDEL) at C-terminal of both prey and bait complex, and using 41AA as Linker1. D, the dynamical range of the YST-PPI system after optimized under the condition of using 2G4S as both Linker1 and Linker2.

Next, we explored the different linker sequences between the target proteins and split-fast TEV fragments to increase the sensitivity of YST-PPI system (**Figure 1A, Figure S4B**). In the original split-TEV complementation system developed in PC12 and COS1 cells ^[17]^, a 41-amino acid polypeptide linker (41AA: ARGRTPPSLGPQDESCTTASSSLAKDTSSASPSNPGASNGS) was utilized between the target protein and N-fast TEV, which consisted of a 12-amino acid semi-flexible sequence (AS: ASPSNPGASNGS) ^[24]^ and the C-terminal tail of the arginine vasopressin receptor 2 to enhance the complex sensitivity and stability ^[25]^ in mammalian cells. Under the condition of the weak ERS (HDEL) at C-terminal of both prey and bait complex, and 2G4S Linker2, the original 41AA Linker1 showed a Yae1p-Lto1p interacting score of 20% (**Figure 2B**). In comparison, the different AS, 2G4S, 3G4S, and 4G4S Linker1 sequences presented the Yae1p-Lto1p interacting score of 35%, 60%, 54%, and 53%, respectively (**Figure 2B**). Similarly, the original 2G4S Linker2 also showed the highest Yae1p-Lto1p interacting score of 63% (**Figure 2C**) under the condition of using 41AA as Linker1 (**Figure 1A**). The 3G4S and 4G4S Linker2 sequences presented a slightly lower Yae1p-Lto1p interacting score of 58% and 56%, respectively (**Figure 2C**).

With all these adjustable parameters, we finally characterized the dynamical range of the YST-PPI system by combining different ERS, transcriptional promoters, and linkers. When prey/bait protein complexes and substrate complex have no fused ERS at the C-terminal (**Figure 2D**, None), the original GAL1-GAL10 bidirectional promoter was used, and both Linker1 and Linker2 were kept as 2G4S, the interacting score of Yae1p-Lto1p was 15% (**Figure 2D**, None). Comparably, after ERS was applied to the YST-PPI system, such as KDEL, HDEL, TVHDEL, EHDEL, FEHDEL, and WEHDEL, the Yae1p-Lto1p interacting scores were increased to 25%, 60%, 80%, 90%, 95%, and 96% (**Figure 2D, Figure S4D**), respectively. More importantly, it is noticed that under the same condition of 96% Yae1p-Lto1p interacting score, the Yae1p-Rli1p interacting score was still only about 4% (**Figure 2D**), suggesting that the background the YST-PPI system is very low.

### 3.3 Interactions of ER resident proteins in yeast ER

Genomic analysis reveals that there are 15 proteins identified to contain the -DEL ERS at their C-terminus in *S. cerevisiae*, including Lhs1p, Kar2p, Pdi1p, Cpr5p, Eug1p, Scj1p, Kre5p, Mpd1p, Mpd2p, Sil1p, Sec20p, Sed4p, Gpi17p, Yos9p, and Qcr2p (**Table S1**). Previous studies indicated that most of them are ER resident proteins, except Qcr2p, which contains -PYLDEL sequence at C-terminus and is reported to be located in mitochondrion ^[26]^. Considering the low background and high sensitivity, we chose the WEHDEL condition in **Figure 2D** to characterize the interactions between these 15 resident proteins. In addition, considering that fusion of target proteins to either N-fast TEV or C-fast TEV fragments might affect the final assembly of TEV protease, two formats of construct modules, Y1 and Y2, were generated (**Figure 3A**), in which all 15 selected proteins were subcloned into both Y1 and Y2 constructs by being either preys or baits. This finally led to generation of total 210 constructs and two sets of interacting scores (dataset-Y1 (**Figure S5**) and dataset-Y2 (**Figure S6**)), representing the target protein to be fused with either N-fast TEV or C-fast TEV fragments (**Figure 3B**).

**FIGURE 3.**
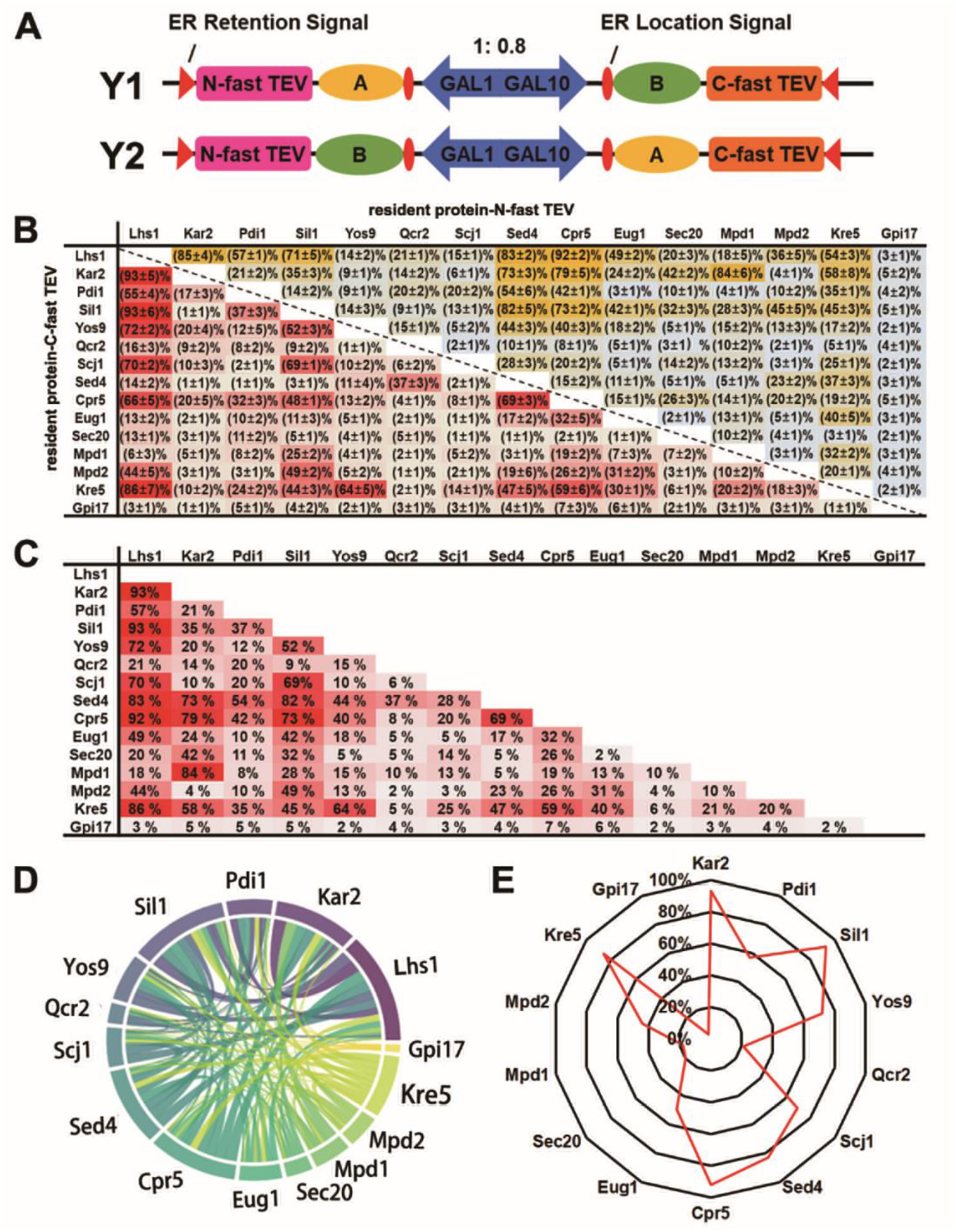
Interaction of ER resident proteins in yeast ER. A, the different patterns of fusion expression of target proteins with N-fast TEV or C-fast TEV fragments. X and Y represent each of the ER resident proteins, respectively. B, the heap-map diagram of the interacting intensity between ER resident proteins. The ER resident proteins are fusion with N-fast TEV fragment as prey (horizontal axis), and C-fast TEV fragment as bait (vertical axis), respectively. The data in the table is the normalized interacting score. Three independent experiments were performed for statistical analysis. C, the final heat-map diagram of the interacting scores between ER resident proteins by data consolidation and filtering of databases Y1 and Y2. D, a circular visualization of protein interaction databases. Different colors are used to distinguish between different ER resident proteins. The thickness of the line indicates the interacting score of the two resident proteins, and the size of the arc indicates the sum of the strength of the interaction between the resident protein and all other proteins. E, the Radar map of Lhs1p interaction intensity with the other 14 resident proteins.

Our results showed that the interacting scores of total 210 protein pairs were in the range of 1% to 93% (**Figure 3B**), which matched the interacting score range of the negative control (Yae1p-Rli1, 4%) and positive control (Yae1p-Lto1, 96%). As expected, the Y1 and Y2 formats did give different interacting scores for the same pair of target proteins. For example, the interacting score of Kar2p-C-fast TEV/Sed4p-N-fast TEV is 73%, indicating a moderate to strong interaction, while the interacting score of Sed4p-C-fast TEV/Kar2p-N-fast TEV is 1%, indicating a non-interaction. We speculated that the N-fast TEV or C-fast TEV fragment might cause the structural hindrance to specific target protein, thus affecting its interaction with other proteins. Considering the low non-specific binding background in the YST-PPI system, we believe the higher interacting score of a specific protein pair in dataset-Y1 and dataset-Y2 is closer to its real interacting capability. For example, the final interacting score of Kar2p/Sed4p is 73%. Therefore, we combine the dataset-Y1 and dataset-Y2 by keeping the higher interacting scores of the specific protein pairs (**Figure 3C**). A total of 91 pairs of ER resident proteins had a greater interacting score than the negative control (Yae1-Rli1, 4%), among which 74 protein pairs presented the interacting scores higher than 10%.

Using the YST-PPI system, the interacting scores between the 15 selected proteins are in the range of 2% - 93% (**Figure 3C**), in which Lhs1p and Gpi17p presented the most and fewest interactions with other proteins, respectively (**Figure 3D**). It was noticed that Lhs1p showed moderate to strong interactions with most of other proteins (interacting score > 40%), except Qcr2p (22%), Sec20p (20%), Mpd1p (18%), and Gpi17p (2%) (**Figure 3E**). Besides Lhs1p, Kar2p, Sil1p, Cpr5p, Kre5p and Sed4p also showed good interactions with most of other target proteins. Another interesting finding is that Lhs1p showed strong interactions with these 5 target proteins, having the interacting scores from 83%-93%. Comparably, all the interacting scores of Gpi17p with other target proteins are lower than 7%. Similar to Gpi17p, Qcr2p and Sec20p also presented low interaction with other target proteins (**Figure 3D**).

Gpi17p is a type III membrane protein, which contains two transmembrane structures ^[27]^. No physical interaction between Gpi17p and the other 14 resident proteins has been reported. However, we cannot exclude the possibility that its structure might be disrupted by N-fast TEV or C-fast TEV fragments. Besides, its membrane location might also lower its interaction with other proteins. Similarly, Sec20p might have the same complexity to cause its low interacting with other proteins. The C-fast TEV fragment fused Sec20p showed low interaction with other target proteins, but the N-fast TEV fragment fused Sec20p presented moderate interaction with Kar2p (42%), Sil1p (32%), and Cpr5p (26%).

### 3.4 The effects of Lhs1 expression levels on ER resident protein interactions

Our results indicated that Lhs1p had the interactions with most of other ER resident proteins (**Figure 3E**), especially strong interactions with several important ER resident proteins, such as Kar2p, etc. We speculate that Lhs1p might play a key regulatory role in the mutual interacting network of ER resident proteins. To explore its potential regulatory roles, *lhs1* knock-out strain EBY100(Δ*lhs1*) (**Figure S1**) and overexpressing strain EBY100(+*lhs1*) (**Figure S2**) were generated, in which interactions of 14 selected protein pairs were characterized and compared with those in the non-modified EBY100 strain (**Table 1, Figure S7**). Our results showed that the interactions between Sil1p and other resident proteins are not changed in either EBY100(Δ*lhs1*) or EBY100(+*lhs1*) strains. However, the interaction scores of other pairs of resident proteins decreased significantly in EBY100(Δ*lhs1*) strain. For example, the interaction scores of Kar2p-Sed4p, Kar2p-Cpr5p, Kar2p-Mpd1p, Pdi1p-Cpr5p, Kre5p-Pdi1p, Kre5p-Cpr5p, Kre5p-Yos9p, Kre5p-Mpd1p decreased from 72%, 78%, 86%, 32%, 23%, 59%, 64%, 21% to 34%, 31%, 65%, 18%, 9%, 22%, 16%, 8%, respectively (**Table 1**). Their interactions in EBY100(+*lhs1*) strain recovered back to the similar levels of those in EBY100 strain. These results indicated that the expression level of Lhs1p could regulate the interactions of the ER resident proteins except the interactions of Sil1p to other proteins.

**TABLE 1:** The effect of *lhs1* expression level on protein synergistic interaction.

| protein1 | protein2 | EEBY100 | EBY100($\Delta lhs1$) | EBY100(+*lhs1*) |
| --- | --- | --- | --- | --- |
| Sed4 | Kar2 | (72±3) % | (34±5) % | (72±6) % |
| Cpr5 | Kar2 | (78±5) % | (31±6) % | (70±5) % |
| Mpd1 | Kar2 | (86±2) % | (65±3) % | (80±4) % |
| Pdi1 | Cpr5 | (32±3) % | (18±1) % | (32±6) % |
| Cpr5 | Sil1 | (73±3) % | (73±5) % | (69±2) % |
| Scj1 | Sil1 | (69±2) % | (65±2) % | (69±5) % |
| Sed4 | Sil1 | (87±6) % | (90±4) % | (82±3) % |
| Yos9 | Sil1 | (52±3) % | (55±2) % | (52±3) % |
| Pdi1 | Sil1 | (37±3) % | (32±6) % | (34±3) % |
| Kre5 | Sil1 | (44±3) % | (38±5) % | (40±5) % |
| Pdi1 | Kre5 | (23±1) % | (9±1) % | (27±2) % |
| Cpr5 | Kre5 | (59±6) % | (22±2) % | (57±3) % |
| Yos9 | Kre5 | (64±5) % | (16±4) % | (63±2) % |
| Mpd1 | Kre5 | (21±3) % | (8±1) % | (20±2) % |

## 4 DISCUSSION

Studying protein interaction can provide guidance of understanding protein’s physiological functions. Inside the ER, the ER resident proteins, which most are molecular chaperones, play key roles in maintaining ER’s function. It is reasonable to believe that these ER resident proteins might interact each other to carry out their functions. However, none of the existing methods is located into the ER lumen for in situ PPI characterization. For example, the interactions between Erd1/Erd2 and the ERS of HDEL, FEHDEL, and WEHDEL, which specifically happen in ER, could not be detected using yeast two-hybrid (Y2H) in our previous studies ^[18]^. Wang et al. characterized genome-wide CCIs (crimp-coil interaction) networks in *S. cerevisiae* using a classical yeast two-hybrid (Y2H) system ^[28]^, in which the interaction between Lhs1p and Kar2p was identified. Nevertheless, the Y2H is anchored in the nucleus, the interactions could not reflect their physiological conditions in yeast ER. Similar concerns with other reported researches, such as using the tandem affinity purification combined with machine learning scoring ^[29]^ and evaluating the genetic interaction using gene mutated yeast strains ^[30]^ to characterize the whole cell PPIs in *S. cerevisiae*, that they could generate comprehensive databases but not specifically focus on analyzing the PPIs in yeast ER.

Some other techniques, such as split fluorescent proteins, split enzymes (luciferase) might provide the basis for the development of new techniques for intracellular detection of protein interactions, including split-GFP/YFP ^[31, 32]^, split-luciferase ^[33]^ and other technologies. However, split-GFP/YFP technology only has fluorescence amplification effect, and the stability of fluorescence signal is not strong enough ^[34, 35]^. The splits-luciferase complementation assays have a two-stage amplification effect to detect weak PPIs, but requiring additional fluorescein substrates and the reaction is susceptible to inhibitors and solutes ^[36]^. Comparably, the YST-PPI system developed here could quantitatively characterize the PPIs in yeast ER in a fast and easy manner. The YST-PPI system builds three cascades amplification effect through ERS, enzymatic reaction, and fluorescence signals to provide a broad dynamic range for characterizing PPIs in ER. The retention effect of ERS ensures that the two target protein complexes have enough time and opportunity to interact in the ER, enabling the primary amplification of the signal. Subsequently, the target protein interaction activated the TEV-fast protease to convert the protein interaction intensities into substrate cleavage, achieving the stable preservation and secondary amplification of the interaction information. Finally, the fluorescence labeling of the displayed substrate converts the enzymatic outcomes into fluorescent signals, which can be quantitatively analyzed by flow cytometry. This three-cascade amplification makes the YST-PPI system highly sensitive and at the same time retains qualitative and quantitative information about instantaneous interactions, providing a new method for detecting the true interactions of proteins in the ER under physiological conditions.

We also tried to regulate the sensitivity of YST-PPI system through different transcriptional promoters to adjust the target proteins concentrations in yeast ER. Four promoter sequences, G4BS13, G4BS3, G4BS4, and G4BS24 (**Figure S8A**), which showed transcriptional strength of ½, ⅓, ¼ and ⅛ of the wild-type UAS_GAL_, respectively ^[37]^, were evaluated by replacing the UAS_GAL_ of bidirectional GAL1-GAL10 promoter in YST-PPI system (**Figure S8B, S8C**). Under the condition of using WEHDEL to evaluate the interaction of Yae1p-Lto1p, those recombinant promoters presented normalized interacting scores of 80%, 75%, 68%, and 63%, respectively (**Figure S8B**). Among them, the transcriptional strength of wild-type UAS_GAL_ was eight times higher than that of G4BS24, while the interacting score of Yae1p-Lto1p only decreased from 90% to 63% (**Figure S8B**). These results indicated that different promoters can further regulate the sensitivity of the YST-PPI system, while the ERS gives the strongest regulatory effects.

According to searching in the most updated protein interaction database BioGRID ^[11]^ and STRING ^[12]^, 24 out of total 105 protein pairs (Lhs1p-Kar2p, Lhs1p-Pdi1p, Lhs1p-Scj1p, Lhs1p-Eug1p, Kar2p-Pdi1p, Kar2p-Sil1p, Kar2p-Yos9p, Kar2p-Scj1p, Kar2p-Eug1p, Pdi1p-Sil1p, Pdi1p-Yos9p, Pdi1p-Scj1p, Pdi1p-Cpr5p, Pdi1p-Eug1p, Pdi1p-Mpd1p, Pdi1p-Mpd2p, Pdi1p-Kre5p, Eug1p-Sil1p, Eug1p-Yos9p, Eug1p-Scj1p, Eug1p-Cpr5p, Eug1p-Mpd1p, Eug1p-Mpd2p, Eug1p-Kre5) have been reported to be interacted based on *in vitro* experiments. Most of these PPIs are also confirmed in our studies with the interesting scores in the range of 5% to – 93%. BioGRID is a biomedical interaction repository with data compiled through comprehensive curation efforts, including physical and genetic interactions between proteins or genes. The interaction in STRING database contains the direct physical interaction and indirect functional correlation between proteins, deriving from experimental data as well as bioinformatics predictions. Comparably, the results obtained here confirm their physiological interactions in the yeast ER, and meanwhile provide the quantitative interacting information. There are also several protein pairs showing low interesting scores in the YST-PPI system, such as Eug1p-Scj1p (5%), and Kar2p-Scj1p (10%). Considering the target proteins are ER resident proteins, the PPIs characterized in the YST-PPI system should be closer to their physiological status.

In the YST-PPI system, the interacting scores also bring valuable speculations of the physiological functions of target proteins. For example, Lhs1p and Sil1p are two important and interesting ER resident proteins, and we speculate that Lhs1p and Sil1p have complementary functions and they interact either directly or through other bridging proteins based on our studies. First, both Lhs1p and Sil1p interact with most of other tested proteins with Lhs1p having higher interacting scores (**Figure 3C**). Secondly, the interactions of Sil1p with other proteins are not affected in the EBY100(Δ*lhs1*) strain (**Table 1**). Moreover, it was reported that double deletion (Δ*lhs1*Δ*sil1*) is lethal while single deletions of *LHS1* (Δ*lhs1*) or *SIL1* (Δ*sil1*) in yeast cells are viable ^[38, 39]^. These results indicate that Lhs1p and Sil1p might have complementary functions in cells, and Sil1p helps rescue the cell viability when Lhs1p is malfunctional. In fact, it was reported that chaperone activity of Kar2p can be activated by both Lhs1p and Sil1p ^[40]^. The interaction between Lhs1p and Kar2p is more precisely controlled by requiring binding to both the nucleotide binding region and linker region, while Sil1p can stimulate Kar2p through just binding to its IIB region. In addition, the mutation of highly conserved residues within the IIB domain inhibits the ability of Sil1p to stimulate the ATPase activity of Kar2p, but it fails to perturb Lhs1p activating Kar2p ^[40]^. Interestingly, the YST-PPI system detected the interaction strength of Kar2p with Lhs1p and Sil1p at 93% and 35% (**Figure 3C**), respectively, which might suggest Kar2p-Lhs1p is the main activation route in cells. One intriguing funding is that our YST-PPI system detected a strong interaction between Lhs1p and Sil1p with an interacting score of 93%, while it was previously reported that no interaction can be detected between these two proteins under *in vitro* characterization ^[41]^. This might suggest Lhs1p interacts with Sil1p either directly or through other bridging proteins, but if it plays a regulatory role of Sil1p in cells still needs to be further explored.

It is very possible that the molecular chaperones in ER could form multimer complex to carry out their functions, which is also indicated in our studies. For example, previous research showed that Kar2p interacted with Yos9p-Hrd3p complex to stabilize it on the E3 complex ^[42, 43]^. In YST-PPI system, the interaction score of Kar2p-Yos9p is only 20%, suggesting that their strong interaction might be mediated by other ER proteins. In addition, it was also reported that Scj1p, Lhs1p and Sil1p all control the binding or release of Kar2p against substrate polypeptides by regulating its ATP hydrolysis ^[40]^, whose interaction with Kar2p was also confirmed by the YST-PPI system. Interestingly, the interaction scores among Scj1p, Lhs1p and Sil1p were all high, ranging from 69% to 93%. It is very possible for them to form multimer complex with Kar2p, which needs to be further verified.

## AUTHOR CONTRIBUTIONS

Xian Fan, Huahua He, Ting Wang, Pan Xu, Faying Zhang, Shantong Hu, Meng Mei, and Yueli Yun conducted the experiments; Li Yi and Meng Mei contributed to the conceptualization; Xian Fan analyzed the data and drafted the manuscript; Li Yi, and Guimin zhang conceived the idea for the project and wrote the paper with Xian Fan, and Meng Mei. All authors have read and agreed to the published version of the manuscript.

## ACKNOWLEDGMENTS

This word was supported by the National Key Research and Development Program of China (NO.2018YFA0901100 to L. Yi and G. Zhang), the National Natural Science Foundation of China (No.32000032 to M. Mei and No. 32370101 to G. Zhang). L.Y. and G.Z. also acknowledge the support from State Key Laboratory of Biocatalysis and Enzyme Engineering.

## DATA AVAILABILITY STATEMENT

The data that support the findings of this study are available from the corresponding author upon reasonable request.

