## Supplementary material for "Characterizing interactions of ER resident proteins in situ through the YST-PPI method": 1

**Running title**: *Quantitative protein interaction characterization in yeast ER*

Xian Fan**^#^**^1^, Huahua He**^#^**^1^, Ting Wang^1^, Pan Xu^1^, Faying Zhang^1,2^, Shantong Hu^2^, Yueli Yun^1^, Meng Mei^1^, Guimin Zhang*^1, 2^, Li Yi*^1^

^1^ State Key Laboratory of Biocatalysis and Enzyme Engineering, Hubei Collaborative Innovation Center for Green Transformation of Bio-resources, Hubei Key Laboratory of Industrial Biotechnology, School of Life Sciences, Hubei University, Wuhan, 430062, China

^2^ College of Life Science and Technology, Beijing University of Chemical Technology, Beijing 100029, China.

^#^ Contributed equally.

**Key word**: YST-PPI; protein-protein interaction; in situ detection; ER resident protein; *S. cerevisiae*;

**Supporting Figures**

**Figure S1**

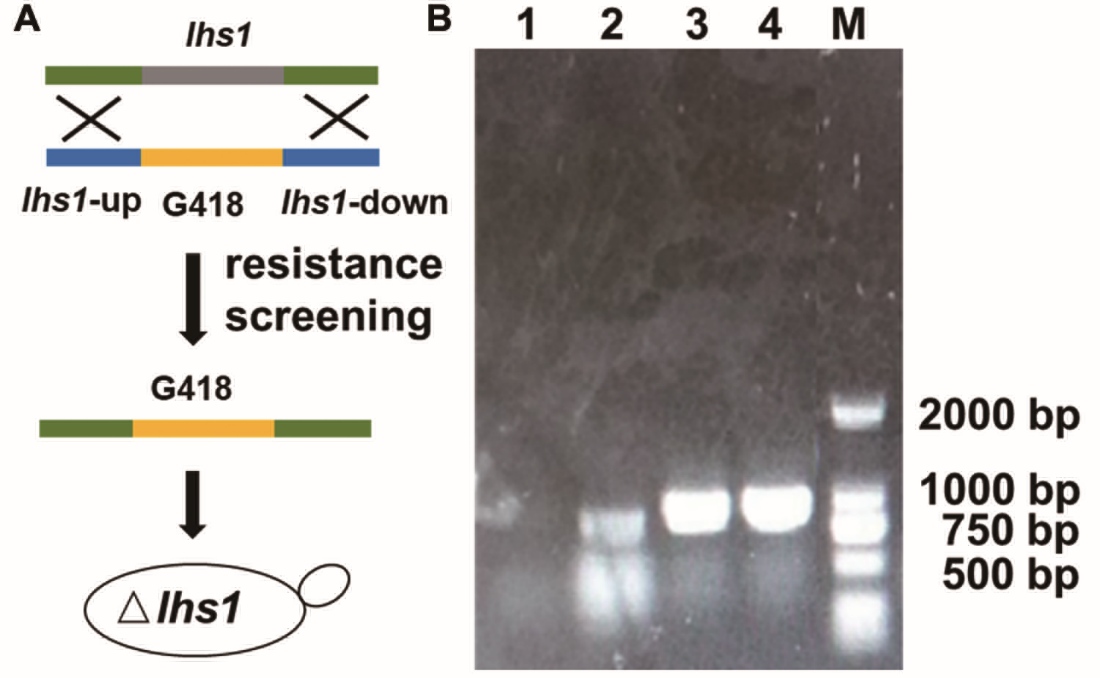

**FIGURE S1. The construction of EBY100(Δ*lhs1*) strain.** A, the schematic diagram of EBY100(Δ*lhs1*) strain construction process. The in EBY100, the *lhs1* gene fragment is replaced by the *G418* gene fragment to generate the EBY100(Δ*lhs1*) strain. B, the agarose gel diagram of EBY100(Δ*lhs1*) strain validation. In EBY100(Δ*lhs1*) strain, the *G418* gene band (line 3, 809 bp) was amplified, but the *lhs1* gene band (line 1) was not obtained. The upstream homologous arm fragment of *lhs1* gene (line 2) and the downstream homologous arm fragment of *lhs1* gene (line 4) was also in line with expectations. Lane M is the DNA ladder.

**Figure S2**

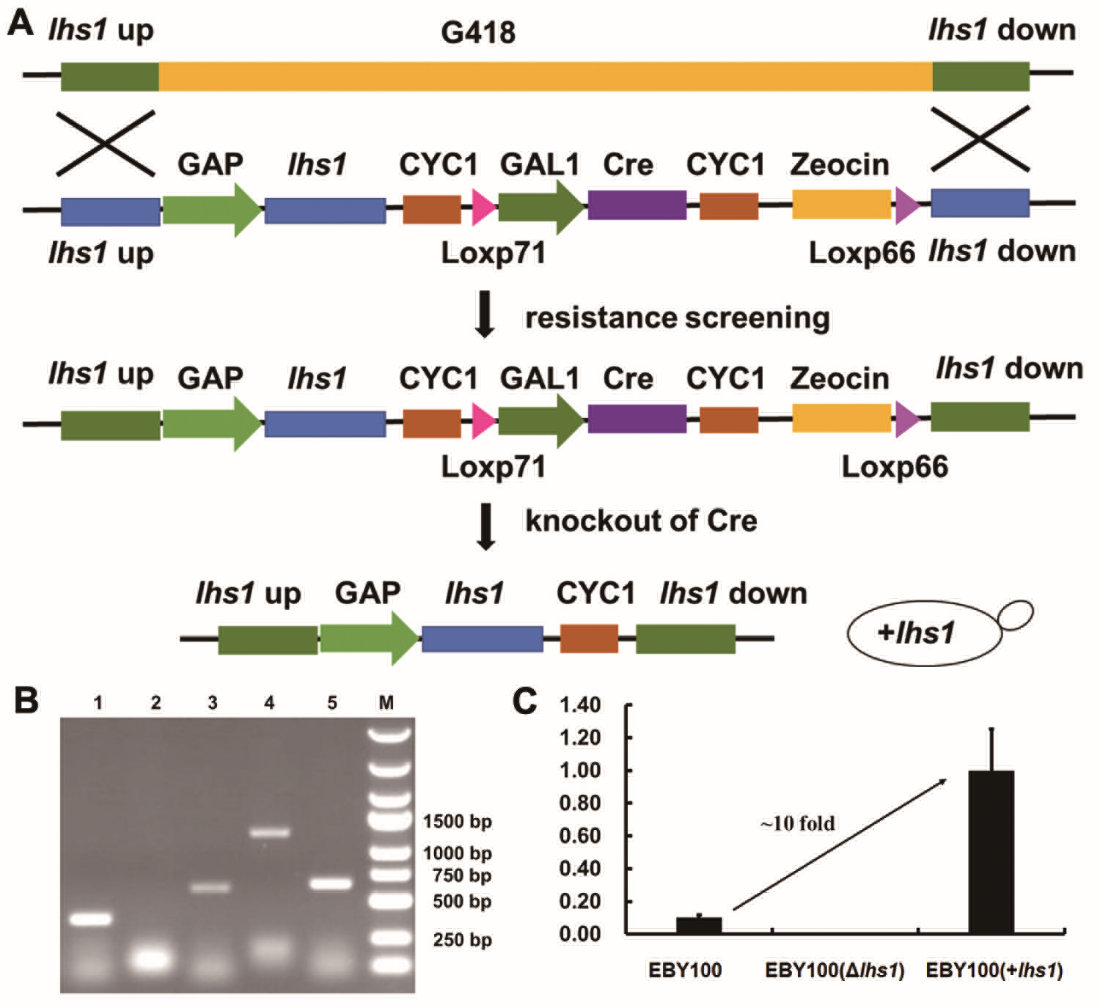

**FIGURE S2. The construction of EBY100(+*lhs1*) strain.** A, the schematic diagram of EBY100(+*lhs1*) strain construction process. In the EBY100(Δ*lhs1*) strain, the *G418* gene fragment is replaced by the GAP-*lhs1* gene fragment to generate the EBY100(+*lhs1*) strain. The GAP strong promoter will lead to the overexpression of Lhs1. B, the DNA agarose gel diagram of EBY100(+*lhs1*) strain validation. The strains that grew on the *Zeocin* resistant plate but not on the *G418* resistant plate were selected as successfully integrated strains, and the *Zeocin* resistance gene (line 1, 372 bp) but not the *G418* gene (line 2, 809 bp) was amplified in the integrated cells. In order to avoid the influence of *Zeocin* resistance gene on integrated strains, we induce Cre expression to knock out the gene fragment (-GAL1-Cre-CYC1-*Zeocin* -). The final integrated strain (EBY100(+*lhs1*)) that did not carry the resistance gene was amplified and verified. The partial fragment of *lhs1* genes (lane 3), the upstream homologous arm fragment of *lhs1* gene (lane 4) and the downstream homologous arm fragment of *lhs1* gene (lane 5) were all in line with expectations. Lane M is the DNA ladder. C, the transcription levels of *lhs1* gene in EBY100, EBY100(Δ*lhs1*) and EBY100(+*lhs1*) strains, respectively.

**Figure S3**

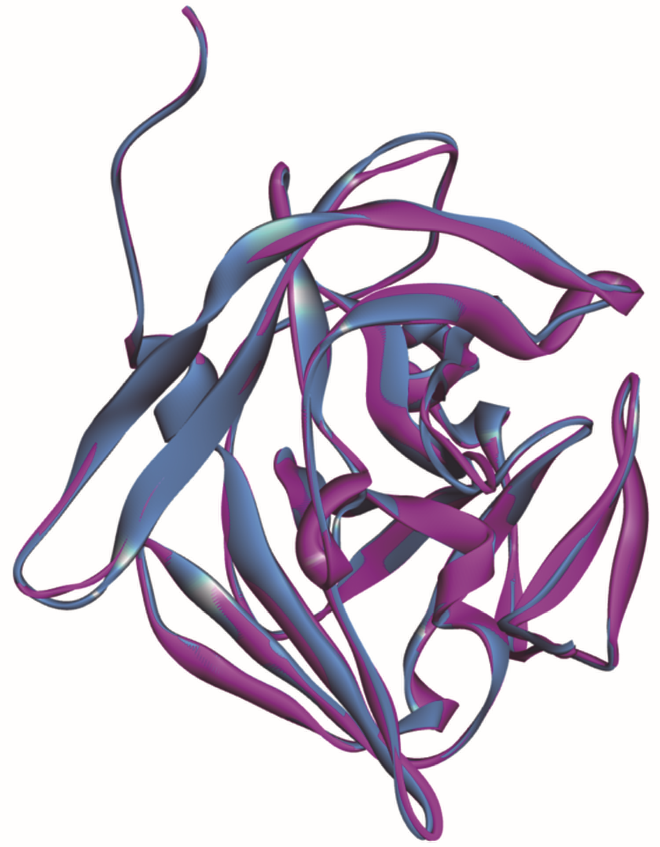

**FIGURE S3. The structure alignment of TEVp and fast TEVp.** The red is TEVp (1LVB^[1]^) and the blue is fast TEVp. The fast TEVp structure is simulated by Alphafold2 ^[2]^.

**Figure S4**

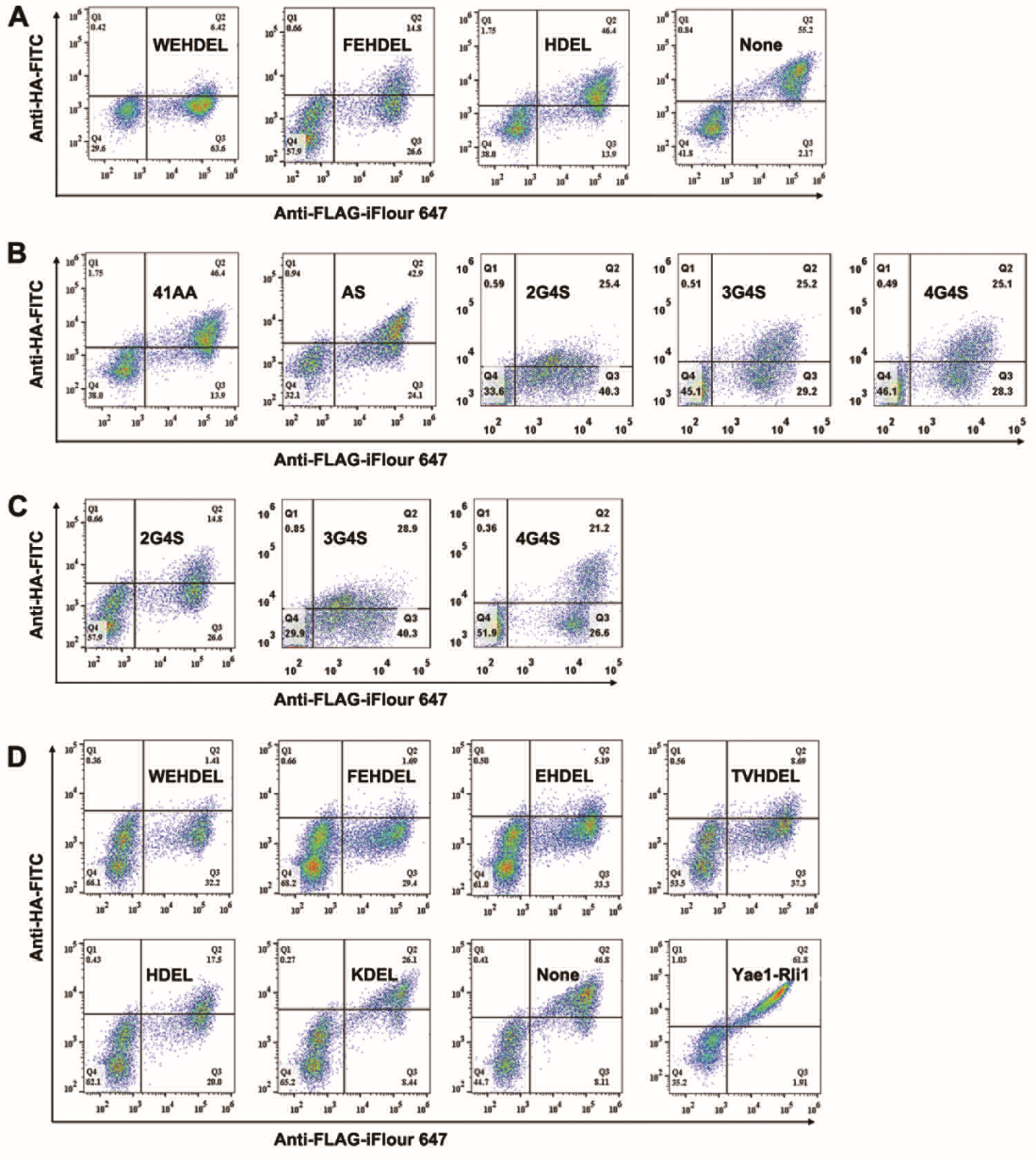

**FIGURE S4. Quantitative analysis of the sensitivity of YST-PPI system.** The Yae1p and Lto1p was fused with N-fast TEV and C-fast TEV, respectively. The Yae1p-Lto1p interacting score was calculated to evaluate the sensitivity of YST-PPI system. A, the effect of different ERS under the condition of using 41AA as Linker1 and 2G4S as Linker2. B, the effect of different Linker1 under the condition of the weak ERS (HDEL) at C-terminal of both prey and bait complex, and using 2G4S as Linker2. C, the effect of different Linker2 under the condition of the ERS (FEHDEL) at C-terminal of both prey and bait complex, and using 41AA as Linker1. D, the dynamical range of the YST-PPI system after optimized under the condition of using 2G4S as both Linker1 and Linker2.

**Figure S5**

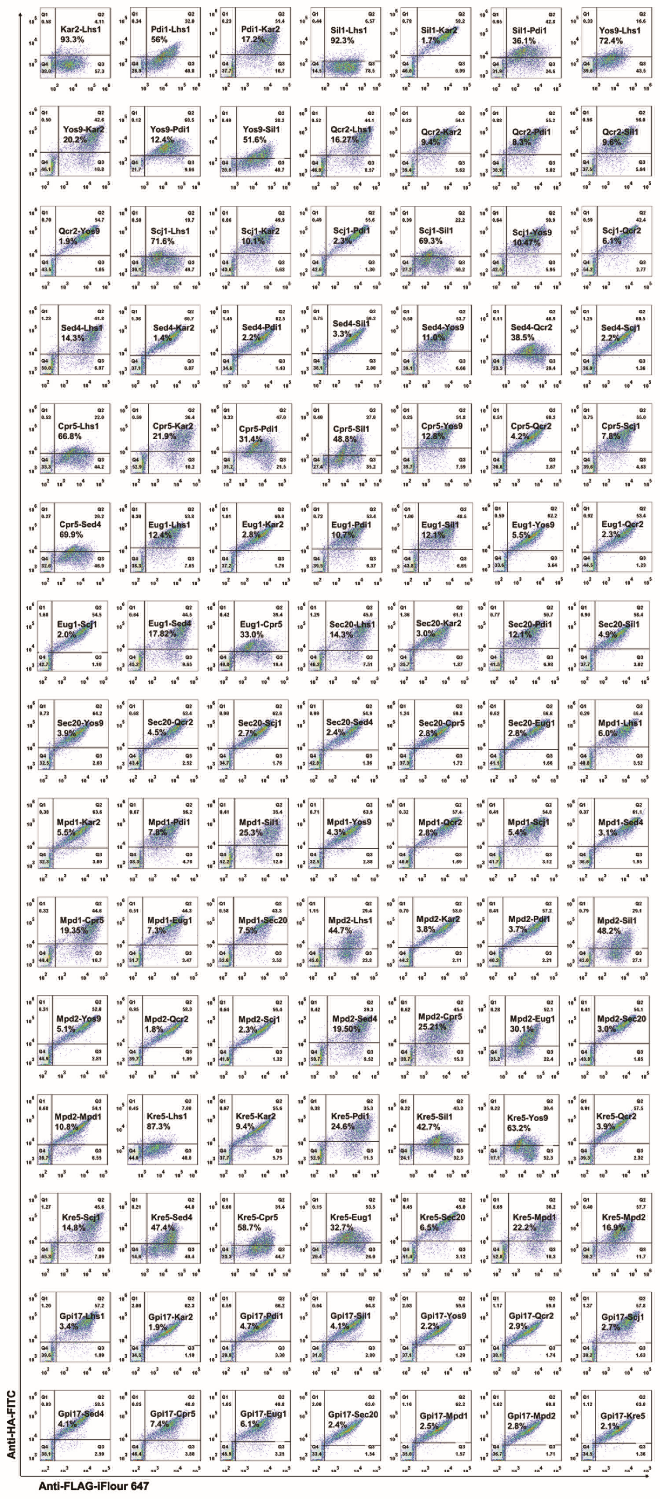

**FIGURE S5. Representative quantitative analysis of ER resident proteins** **interactions by Y1 mode.**

**Figure S6**

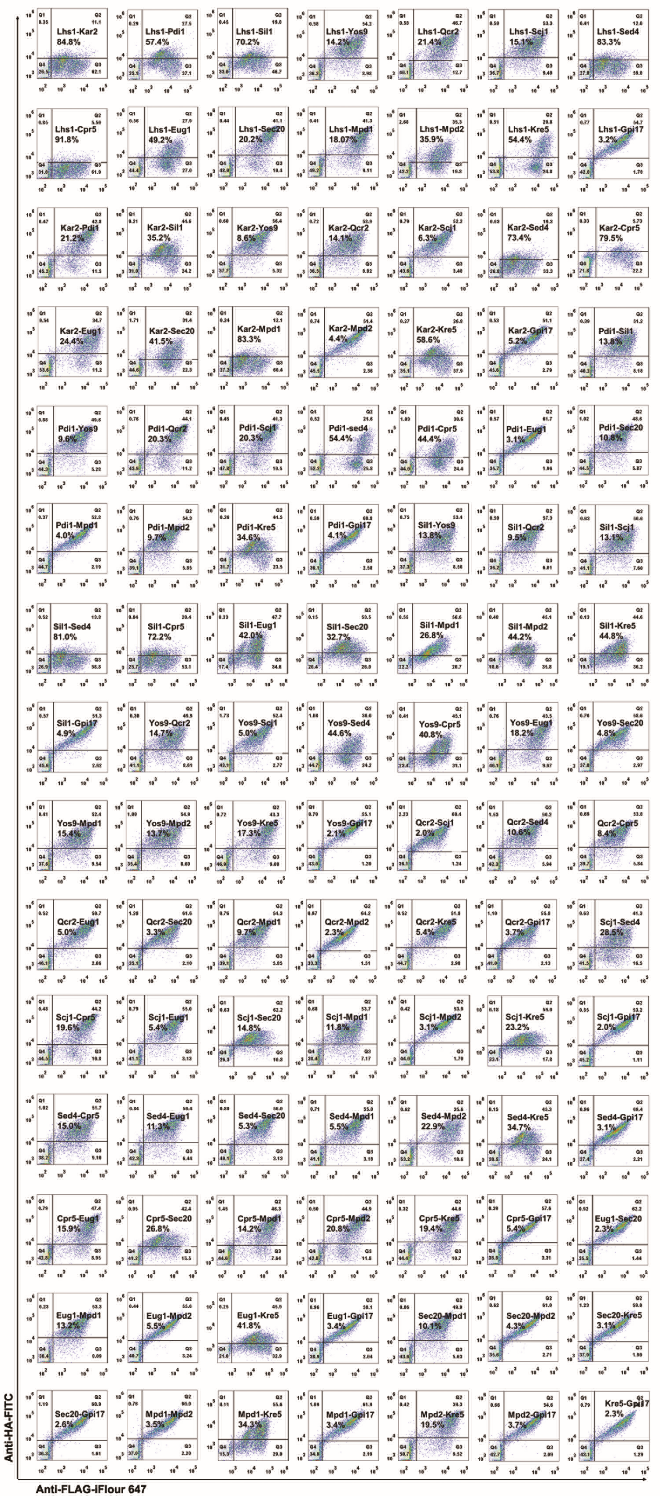

**FIGURE S6. Representative quantitative analysis of ER resident proteins** **interactions by Y2 mode.**

**Figure S7**

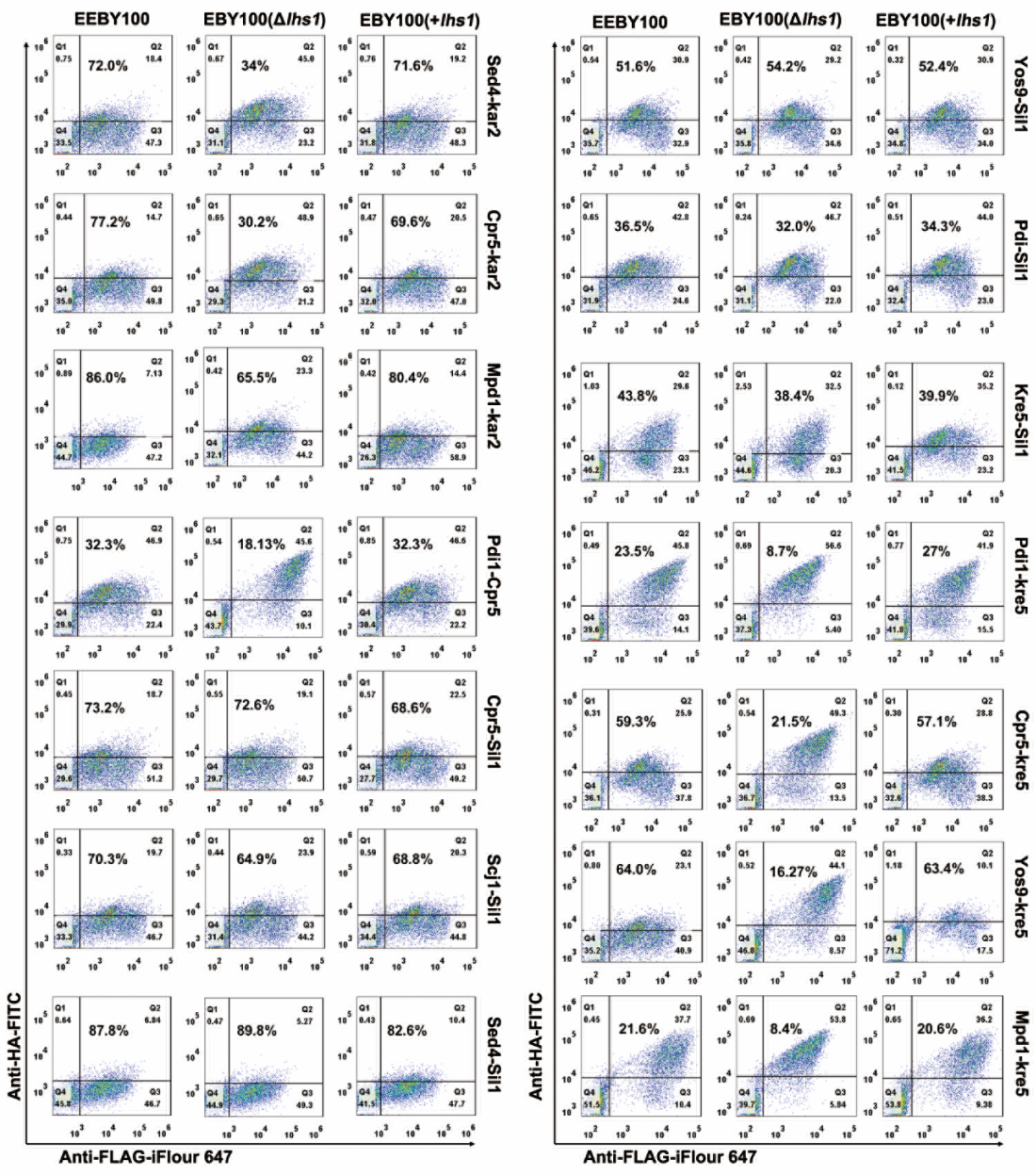

**FIGURE S7. Representative quantitative analysis of the effect of *lhs1* expression level on selected protein-protein interaction.**

**Figure S8**

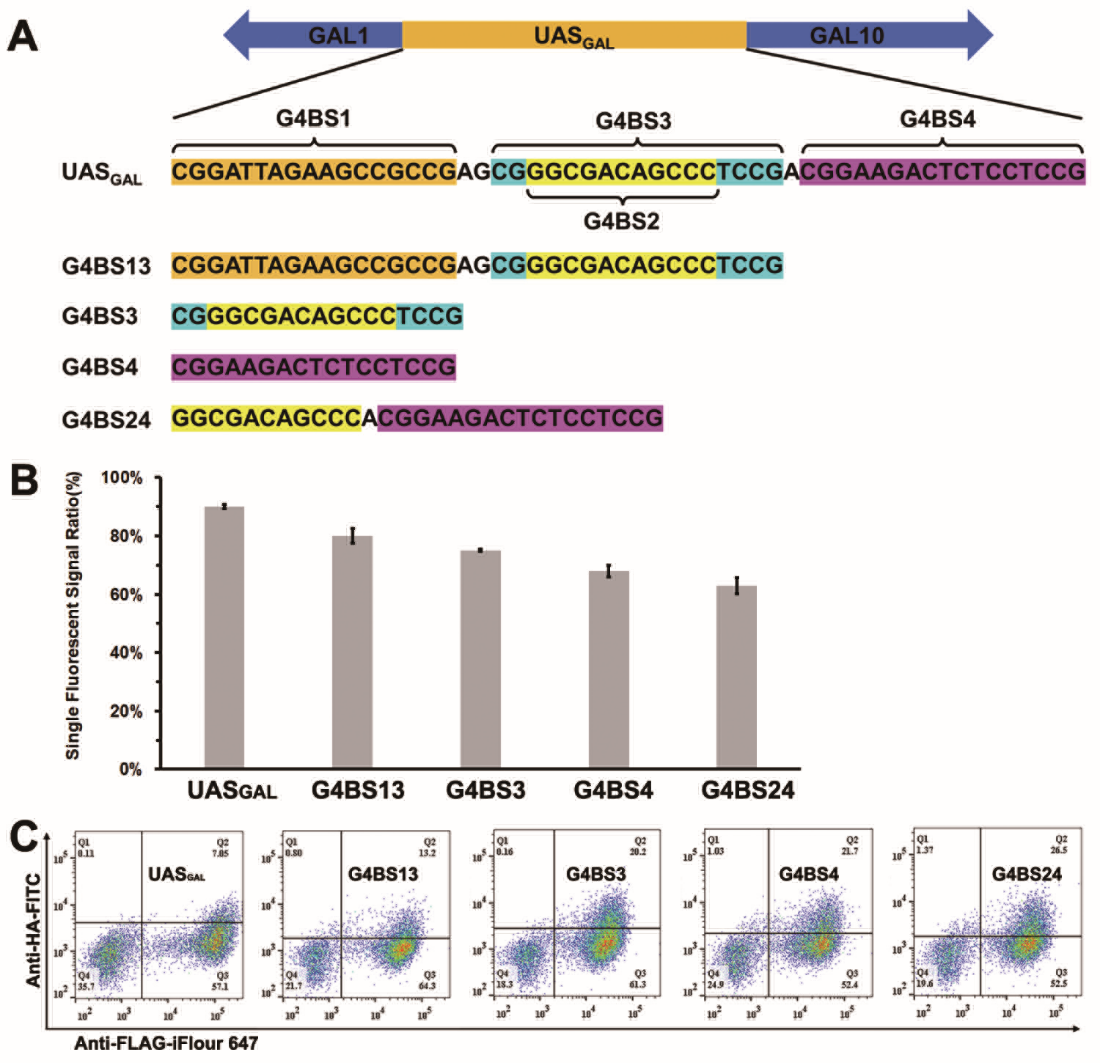

**FIGURE S8. The reconstructed UAS_GAL_ sequences.** A, the G4BS1, G4BS2, G4BS3, and G4BS4 sequence are marked in orange, yellow, blue and yellow, rose red, respectively. The UAS_GAL_ of bidirectional GAL1-GAL10 promoter in YST-PPI system was replaced by reconstructed G4SB13, G4SB3, G4SB4 and G4SB24 sequences to systematic optimization. B, the interacting scores of different promoters using fast TEV protease as split protein under the condition of the strongest ERS (WEHDEL) at C-terminal of both prey and bait complex, using 41AA as Linker1 and 2G4S as Linker2. C, the Yas1p/Lto1p interactions under the condition of different promoters.

**Supporting Table**

**TABLE S1. The ER resident protein of yeast and human.**

| **protein** | | **Uniprot** | | **ERS** | | **function** | **ref** |
| --- | --- | --- | --- | --- | --- | --- | --- |
| **S.c** | **Human** | **S.c** | **Human** | **S.c** | **Human** |  |  |
| Lhs1 | HYOU1 | P36016 | Q9Y4L1 | ILHDEL | LKNDEL | chaperone binding | ^[3, 4]^ |
| Kar2 | Bip | P16474 | P11021 | FEHDEL | AEKDEL | unfolded protein binding | ^[5, 6]^ |
| Sil1 | Sil1 | Q08199 | Q9H173 | NFRDEL | LLKELR | nucleotide exchange factor | ^[7, 8]^ |
| Scj1 | HEDJ | P25303 | Q9UBS4 | MLKDEL | NGLQGY | unfolded protein binding | ^[9, 10]^ |
| Cpr5 | Cyp40 | P35176 | Q08752 | AAHDEL | YAKMFA | peptidyl-prolyl cis-trans isomerase activity | ^[11, 12]^ |
| Pdi1 | PdiA1 | P17967 | P07237 | AIHDEL | AVKDEL | protein disulfide isomerase activity | ^[13, 14]^ |
| Eug1 | PdiA4 | P32474 | P13667 | TVHDEL | RTKEEL | response to endoplasmic reticulum stress | ^[15, 16]^ |
| Mpd1 | PdiA6 | Q12404 | Q15084 | NKHDEL | LGKDEL | protein folding | ^[17, 18]^ |
| Mpd2 | PdiA2 | Q99316 | Q13087 | SSHDEL | GSKEEL | disulfide oxidoreductase activity | ^[19, 20]^ |
| Yos9 | OS9 | Q99220 | Q13438 | IEHDEL | LDEFDF | retrograde protein transport | ^[21, 22]^ |
| Kre5 | - | P22023 | - | PLHDEL | - | protein required for beta-1,6 glucan biosynthesis | ^[23]^ |
| Qcr2 | Qcr2 | P07257 | P22695 | PYLDEL | PFVDEL | subunit 2 of ubiquinol cytochrome-c reductase (Complex III) | ^[24, 25]^ |
| Sed4 | DelGEF | P25365 | Q9UGK8 | GLHDEL | SRNGGL | guanyl-nucleotide exchange factor activity | ^[26, 27]^ |
| Sec20 | Sec20 | P28791 | Q12981 | VSHDEL | RLFPFL | SNAP receptor activity | ^[28, 29]^ |
| Gpi17 | PIGS | Q04080 | Q96S52 | DGEDEL | KPEKTD | GPI-anchor transamidase activity | ^[30, 31]^ |
